# De novo assembly of complete *Plasmodium falciparum* isolate genomes using PacBio HiFi sequencing technology

**DOI:** 10.1101/2025.11.19.689036

**Authors:** Prince B. Nyarko, Mathieu Quenu, Marc-Antoine Guery, Camille Cohen, Godfrey Kinyori Wagutu, William L. Hamilton, Sophia T. Girgis, Alex Makunin, Shane A. McCarthy, Mara K. N. Lawniczak, Antoine Claessens

## Abstract

*Plasmodium falciparum* possesses a highly structured genome with extensive sequence diversity concentrated in Variant Surface Antigen (VSA) families. These genes—*var*, *rif*, and *stevor*—play key roles in immune evasion and pathogenesis and are difficult to assemble using short-read sequencing technologies. Here, we applied PacBio HiFi long-read sequencing to generate high-quality *de novo* genome assemblies from 43 *P. falciparum* parasite cultures originating from community cases in The Gambia. Parasites were culture-adapted, cloned by limiting dilution where possible, and sequenced using high molecular weight DNA extracts. Assemblies from single-genotype lineages were constructed using hifiasm, producing complete chromosomal-length scaffolds with high base accuracy without requiring short-read polishing. We recovered full repertoires of *var*, *rif*, and *stevor* genes and classified them into known subgroups. Together, our results demonstrate that PacBio HiFi sequencing enables accurate assembly of complex *P. falciparum* genomes from natural infections. This work provides a valuable genomic resource for future studies of parasite evolution, transmission dynamics, and antigenic diversity, and suggests that VSA repertoires can serve as reliable proxies of genetic relatedness across infections.

## Introduction

Malaria remains a global leading cause of morbidity and mortality, affecting millions and causing hundreds of thousands of deaths annually (*World Malaria Report 2024*, n.d.) The disease is caused by unicellular apicomplexan parasites of the genus *Plasmodium*, with *P. falciparum* responsible for most of the severe cases and fatalities. The first *P. falciparum* genome sequenced was the laboratory strain 3D7 in 2002 (Gardner et al., 2002). Since then, numerous studies have increased the number of *P. falciparum* genome sequences.

Most genome sequencing efforts have relied on Illumina short-read technologies (MalariaGen., 2021), but long-read platforms, such as Oxford Nanopore Technology or PacBio’s single-molecule real-time (SMRT) sequencing have been increasingly applied to *Plasmodium* genomes (Delandre et al., 2024; Holzschuh et al., 2024; Vembar et al., 2016). Historically, the lower base-calling accuracy of long-read platforms necessitated complementary short-read sequencing to achieve assemblies that were both accurate and complete (Higgins et al., 2024). However, advances in PacBio HiFi (High Fidelity) sequencing using circular consensus sequencing (CCS) now promise long and highly accurate reads (Hon et al., 2020; Wenger et al., 2019). While this technology has been used to produce high-quality assemblies for some *Plasmodium* species (*P. yoelii* - Godin et al., 2023), it has not yet been widely applied to *P. falciparum* field isolates.

The genome of *P. falciparum* consists of a conserved ‘core genome’ and a hypervariable non-core genome, the latter harboring large multigene families encoding Variant Surface Antigens (VSAs) (Otto et al., 2018). The three VSA families - *var*, *rif* and *stevor* - are important for pathogenesis and immune evasion and are located in subtelomeric and internal gene clusters (Harrison et al., 2020; Jespersen et al., 2016; Rowe et al., 2009; Sakoguchi et al., 2025). Their sequences are highly polymorphic with each parasite isolate typically having a unique set of VSAs whose genetic diversity can be generated by non-homologous recombination events (Claessens et al., 2014). Their classification into biologically relevant subgroups (e.g., *var* A, B, and C; *rif* A and B) provides insights into parasite biology and virulence. For example, group A *var* genes are implicated in cerebral malaria (Claessens et al., 2012). Analyses of VSA repertoires in field isolates also bear epidemiological relevance, enabling lineage tracking across host infections (Day et al., 2025; Tonkin-Hill et al., 2021). However, assembling these hypervariable regions with short-read data has proven difficult due to repetitive sequences and limited homology between isolates, often resulting in incomplete sets of VSA sequences.

In this study, we apply PacBio HiFi sequencing to generate *de novo* assemblies of *P. falciparum* parasites from The Gambia. Using *in vitro* culture to obtain high-yield high molecular weight (HMW) DNA, we sequenced both clonal lineages (obtained via limiting dilution) and bulk field-adapted populations, resulting in 43 assemblies. We then used bioinformatics tools to assemble and annotate genomes, extract, classify and analyze VSA sequences.

## Methods

### Study design and sample collection

All *P. falciparum* field isolates reported here have been described previously (Collins et al., 2022; Fogang et al., 2024; Guery et al., 2025). Briefly, 5mL venous blood samples were collected from *P. falciparum* positive participants in 2016 and 2017 in The Gambia. Plasma, PBMCs and red blood cells were immediately separated by centrifugation. Red blood cells were cryopreserved with glycerolyte in liquid nitrogen. The study protocol was reviewed and approved by the Gambia Government/MRC Joint Ethics Committee (SCC 1476, SCC 1318, L2015.50) and by the London School of Hygiene & Tropical Medicine Ethics Committee (Ref. 10982). The field studies were also approved by local administrative representatives, the village chiefs. Written informed consent was obtained from participants over 18 years old and from parents/guardians for participants under 18 years. Written assent was obtained from all individuals aged 12–17 years. Approval under the Nagoya Protocol was granted by the National Focal Point of The Gambia.

### Parasite culturing

Thawed parasites were cultured in RPMI-1640 (sigma) supplemented with 25mM HEPES, 2mM L-glutamine, 0.5% Albumax II (sigma), 50µg/L gentamicin (sigma) and 5% AB^+^ normal human serum (NHS) in static conditions. Parasites were cultured in 10mL volumes at 2% haematocrit in a blood gas environment of 90% N_2_, 5% CO_2_ and 5% oxygen (O_2_).

### Cloning by limiting dilution

Once field isolates were culture-adapted, typically after 2 to 3 weeks, clones were isolated by limiting dilution in 96-well plates at a theoretical density of 1 parasite per well (ppw) – a 3ppw was included as positive control – and maintained at 1.25% haematocrit. Plates were kept in a modular chamber with blood gas and incubated at 37°C. On day 7, exhausted media was removed and replaced with fresh media containing uninfected RBCs (uRBCs) at 1.25% haematocrit. Culture media was changed every two days from this point onwards. From day 14, 3ppw wells were screened by microscopy for viable parasites. If parasites were absent, plates were screened every two days until day 21, at which point the cloning assay was considered failed and discarded. If viable parasites were present, the entire plate was screened for parasites in the 1ppw wells. Positive wells were initially transferred into 6-well plates with 100µL uRBCs and 3mL media, and grown until parasitaemia was visible by microscopy (usually 2 to 4 replication cycles). Parasites were then genotyped by MSP1 and MSP2 (see details in supplementary methods, Supplementary Figure 1) to assess the number of parasite genotypes present in cultures. Wells with confirmed unique single clones were selected for expansion in 25mL cultures at 2% haematocrit, while all other wells were terminated. Parasites were then grown to higher parasitaemia (∼10%) and harvested at the schizont stage for high molecular weight DNA extraction.

### High molecular weight DNA extraction and PacBio sequencing

High molecular weight (HMW) DNA was extracted from both cloned parasites and ‘bulk’ populations (parasite cultures directly adapted from field isolates) using the Monarch® HMW DNA Extraction Kit for Tissue (New England Biolabs). The tissue kit was chosen over the blood kit due to the protein separation step being able to remove residual cell debris and haemozoin contaminants. Several modifications were made to the extraction procedure. Briefly, RBCs were initially lysed with RBC Lysis Buffer from the Monarch HMW DNA Extraction Kit for Cells & Blood. Free parasite pellets were then washed with 1X PBS until the supernatant was clear. All subsequent steps followed the tissue kit’s protocol. The agitation speed for lysis was reduced to 1500 rpm and in special cases where parasite material was limited, precipitation enhancer was added prior to binding DNA to the glass beads. DNA was solubilised in 200µL of elution buffer, incubated at 37°C for 1 hour to enhance dissolution, stored at 4°C overnight and then frozen at -20°C until sequencing. Prior to sequencing, DNA was cleaned using SPRI (Solid Phase Reversible Immobilization) beads, sheared with g-TUBE (Covaris) aiming for a size of 10kb and assessed for integrity using FEMTO Pulse (Agilent). Samples were sequenced using the Pacific Biosciences (PacBio) high fidelity (HiFi) circular consensus sequencing technology (Wenger *et al*., 2019) on the Sequel IIe plaform, with up to 10 samples per SMRT cell.

### Genome assemblies and annotation

We assembled genomes from PacBio HiFi reads datasets using Hifiasm (H. Cheng et al., 2021), using the -l0 option to assemble haploid genome. Contigs were split, oriented and arranged to the 3D7 chromosome configuration using RagTag, with default parameters (Alonge et al., 2022). Assemblies coming from polyclonal isolates were left in the post-Hifiasm assembly state, with no re-arrangements to the 3D7 chromosome configuration. Companion (Steinbiss *et al*., 2016) was used to annotate clone and bulk assemblies with the following options: pseudogene detection activated, AUGUSTUS (Stanke et al., 2004) for structural annotation and RATT (Otto et al., 2011) for transfer of highly conserved genes from 3D7. Genes annotated as ‘hypothetical proteins’ were not included in the total number of protein-coding genes per genome.

### VSAs extraction

*Var* genes were retrieved from all assemblies by screening for genes annotated as potentially encoding PfEMP1 by the Companion pipeline and filtering for a nucleotide length higher than 2500bp. Each *var* gene had its different domains annotated by submitting its amino acid sequences to the varDOM server (Rask et al., 2010). Copies of *var2csa* were extracted from the list of *var* genes by screening for genes with DBLpam domains. Upstream group affiliation of the remaining *var* genes was performed as in the method described in Pangilinan et al., 2025. Briefly, a phylogenetic tree was constructed using the 500bp upstream nucleotide sequences of newly assembled *var* genes plus reference upstream sequences previously annotated as group A, B or C (Lavstsen et al., 2003; Rask et al., 2010). Sequences of all *var* genes were aligned using MAFFT v7.520 and a phylogenetic tree was inferred from the alignment using FastTree v2.1.11. Upstream sequences were annotated based on the clade of the reference upstream group. The HMM-based software STRIDE (Zhou et al., 2022) was used to retrieve *rif* and *stevor* from the list of Companion proteins, and further annotate the *rif* into *rif* groups A or *rif* group B.

### Estimation of complexity of infection across bulk parasite populations

To infer whether bulk parasite populations were single- or poly-genotype we looked at average heterozygosity levels over a panel of 72,020 core-genome variable loci. This set of variable loci was generated by mapping PacBio HiFi reads to the reference 3D7 genome using minimap2 v2.28 (Li, 2018). SNP calling and generation of VCF files were performed with GATK v4.1.0.5.1 (Van der Auwera et al., 2020), using the HaplotypeCaller pipeline. Variants were filtered based on their quality using bcftools v1.9 (Danecek et al., 2021) and imported in an R environment with the R package vcfR (Knaus & Grünwald, 2017). Genome assemblies were considered to be originating from a single-genotype population if their average heterozygosity level was lower than 0.02 over the entire set of 72,020 variants, which corresponded to the highest heterozygosity level found in clone genome assemblies.

### Estimation of base-calling error rates in hifiasm assemblies

To estimate base-calling error rates level in newly-generated assemblies, we used two *ex vivo* isolates (i.e. without any culturing), DC01m1 and DC10m2, sequenced with Illumina in a previous study (Guery et al., 2025). Illumina reads were mapped to their respective long read assembly using bwa MEM v0.7.17 (Li & Durbin, 2009). The output sam files were then converted and sorted using samtools v1.10 (Li et al., 2009). GATK v4.1.0.5.1 (Van der Auwera et al., 2020) was used again for SNPs calling and generation of VCF files, using the HaplotypeCaller pipeline. Additionally, nucleotide level accuracies in assemblies were also estimated by comparing the k-mer decomposition of PacBio HiFi assemblies to illumina reads datasets via the software merqury (Rhie et al., 2020). Here we used the quality value (qv) score to assess base-calling accuracy, a metric that is generated by looking at k-mers present in high-quality reads dataset (Illumina) but not in the PacBio assemblies.

### Estimation of pairwise Identity By Descent (IBD) and VSAs sharing amongst samples

To estimate core genome relatedness between parasites genomes we calculated the pairwise IBD values between all genomes. Chromosomal segments are in IBD state if they descend from a recent common ancestor, and a pairwise IBD value indicate the average proportion of two genomes that are in IBD state. In practise, PacBio HiFi reads were mapped to the reference 3D7 genome using minimap2 v2.28 (Li, 2018). SNP calling was performed as described above. From this, VCF files were parsed with the R package vcfR to generate a tab-delimited table containing SNPs position and status for each genome. We used hmmIBD v2.0.0 (Schaffner et al., 2018) to calculate the fraction of the genome in an IBD state, from here referred as pairwise IBD values, between each pair of genomes.

In parallel, we assessed the similarity of VSAs repertoires between each single-genotype genome pair by clustering *var*, *rif*, and *stevor* sequences using CD-HIT (Fu et al., 2012). Nucleotide sequences in different isolate genomes were considered identical if their sequence similarity was higher than 99%. After counting the number of shared VSAs between two genomes we calculated the three normalised pairwise VSA sharing indexes *var* sharing, *rif* sharing and *stevor* sharing using the following formula:

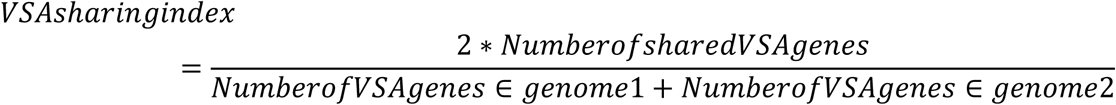

To account for the non-independence of pairwise estimates, overall correlations between core genome IBD values and VSA sharing indexes were then evaluated using Mantel tests, as implemented in the R package vegan (v2.6-10). Distance matrices of pairwise IBD values and normalized *var, rif* and *stevor* sharing were compared via the mante() function, with 9999 permutations and p-values computed using the Spearman rank correlation method.

## Results

Blood isolates were collected in four neighboring villages from The Gambia, following the sample design presented in Collins et al., 2022 and Fogang et al., 2024. Briefly, samples were collected during the wet season (October 2016 – November 2016) from both asymptomatic and symptomatic cases, and from asymptomatic carriers at monthly timepoints during the following dry season (December 2016 to May 2017). From this sampling design, isolates were separated into three infection groups: the Wet season Symptomatics (WS), corresponding to parasites collected from symptomatic cases in the wet season; the Wet season Asymptomatics (WA), corresponding to parasites collected from asymptomatic cases in the wet season and the Dry season asymptomatic Cohort samples (DC), corresponding to parasites collected at monthly timepoints from asymptomatic cases during the dry season. Sample identifiers incorporate group, timepoint, and sequencing type (e.g., bulk ‘b’ vs clone ‘c’; e.g., WA01b, DC01m1c). Metadata and group assignments are detailed in Table 1.

**Table 1:** Genome accession numbers and metadata for 43 *P. falciparum* genome assemblies.

| Genome_ID | Isolate_ID | Group | Cloning dilution | Complexity of infection | Collection date | Host age | Host sex | NCBI accession number |
| --- | --- | --- | --- | --- | --- | --- | --- | --- |
| DC01m1c | DC01m1 | Dry Cohort | clone | single-genotype | 24/12/2016 | 9 | male | ERR9731534 |
| DC02m1c | DC02m1 | Dry Cohort | clone | single-genotype | 24/12/2016 | 11 | male | ERR9731535 |
| DC03m1c | DC03m1 | Dry Cohort | clone | single-genotype | 22/12/2016 | 13 | female | ERR9731539 |
| DC04m2c | DC04m2 | Dry Cohort | clone | single-genotype | 26/01/2017 | 15 | female | ERR9731540 |
| DC07m1b | DC07m1 | Dry Cohort | bulk | single-genotype | 23/12/2016 | 23 | female | ERR12263817 |
| DC07m1c | DC07m1 | Dry Cohort | clone | single-genotype | 23/12/2016 | 23 | female | ERR12263821 |
| DC08m3b | DC08m3 | Dry Cohort | bulk | poly-genotypes | 13/02/2017 | 10 | male | ERR12263802 |
| DC08m3c | DC08m3 | Dry Cohort | clone | single-genotype | 13/02/2017 | 10 | male | ERR12263818 |
| DC08m5b | DC08m5 | Dry Cohort | bulk | poly-genotypes | 19/04/2017 | 10 | male | ERR12263832 |
| DC08m5c1 | DC08m5 | Dry Cohort | clone | single-genotype | 19/04/2017 | 10 | male | ERR12263826 |
| DC08m5c2 | DC08m5 | Dry Cohort | clone | single-genotype | 19/04/2017 | 10 | male | ERR12263801 |
| DC09m5b | DC09m5 | Dry Cohort | bulk | poly-genotypes | 20/04/2017 | 12 | male | ERR12263810 |
| DC09m5c1 | DC09m5 | Dry Cohort | clone | single-genotype | 20/04/2017 | 12 | male | ERR12263806 |
| DC09m5c2 | DC09m5 | Dry Cohort | clone | single-genotype | 20/04/2017 | 12 | male | ERR12263805 |
| DC10m1c | DC10m1 | Dry Cohort | clone | single-genotype | 23/12/2016 | 12 | male | ERR9731536 |
| DC10m2c | DC10m2 | Dry Cohort | clone | single-genotype | 12/01/2017 | 12 | male | ERR9731537 |
| DC10m3c | DC10m3 | Dry Cohort | clone | single-genotype | 12/02/2017 | 12 | male | ERR9731538 |
| WA01b | WA01 | Wet Asymptomatic | bulk | single-genotype | 23/11/2016 | 15 | female | ERR12263816 |
| WA01c | WA01 | Wet Asymptomatic | clone | single-genotype | 23/11/2016 | 15 | female | ERR12263812 |
| WA03b | WA03 | Wet Asymptomatic | bulk | poly-genotypes | 21/11/2016 | 27 | female | ERR12263820 |
| WA03c | WA03 | Wet Asymptomatic | clone | single-genotype | 21/11/2016 | 27 | female | ERR12263808 |
| WA04b | WA04 | Wet Asymptomatic | bulk | single-genotype | 21/11/2016 | 11 | male | ERR12263815 |
| WA04c | WA04 | Wet Asymptomatic | clone | single-genotype | 21/11/2016 | 11 | male | ERR12263823 |
| WA10b | WA10 | Wet Asymptomatic | bulk | single-genotype | 21/11/2016 | 8 | male | ERR12263827 |
| WA10c | WA10 | Wet Asymptomatic | clone | single-genotype | 21/11/2016 | 8 | male | ERR12263807 |
| WA11c | WA11 | Wet Asymptomatic | clone | single-genotype | 23/11/2016 | 9 | male | ERR12263813 |
| WA47b | WA47 | Wet Asymptomatic | bulk | single-genotype | 21/12/2016 | 11 | male | ERR12263803 |
| WA47c | WA47 | Wet Asymptomatic | clone | single-genotype | 21/12/2016 | 11 | male | ERR12263833 |
| WS01b | WS01 | Wet Symptomatic | bulk | poly-genotypes | 08/11/2016 | NA | male | ERR12263825 |
| WS03b | WS03 | Wet Symptomatic | bulk | poly-genotypes | 31/10/2016 | 13 | female | ERR12263830 |
| WS04b | WS04 | Wet Symptomatic | bulk | poly-genotypes | 31/10/2016 | 45 | male | ERR12263837 |
| WS05b | WS05 | Wet Symptomatic | bulk | poly-genotypes | 17/10/2016 | 54 | female | ERR12263811 |
| WS05c | WS05 | Wet Symptomatic | clone | single-genotype | 17/10/2016 | 54 | female | ERR12263828 |
| WS06c | WS06 | Wet Symptomatic | clone | single-genotype | 17/10/2016 | 22 | female | ERR12263800 |
| WS07c | WS07 | Wet Symptomatic | clone | single-genotype | 01/11/2016 | 10 | male | ERR12263814 |
| WS08b | WS08 | Wet Symptomatic | bulk | single-genotype | 07/11/2016 | 41 | male | ERR12263804 |
| WS09b | WS09 | Wet Symptomatic | bulk | single-genotype | 07/11/2016 | 49 | female | ERR12263834 |
| WS10b | WS10 | Wet Symptomatic | bulk | poly-genotypes | 08/11/2016 | 16 | female | ERR12263838 |
| WS11b | WS11 | Wet Symptomatic | bulk | single-genotype | 08/11/2016 | 8 | male | ERR12263824 |
| WS12b | WS12 | Wet Symptomatic | bulk | poly-genotypes | 08/11/2016 | 28 | male | ERR12263831 |
| WS12c | WS12 | Wet Symptomatic | clone | single-genotype | 08/11/2016 | 28 | male | ERR12263822 |
| WS13b | WS13 | Wet Symptomatic | bulk | poly-genotypes | 18/11/2016 | 64 | female | ERR12263836 |
| WS15b | WS15 | Wet Symptomatic | bulk | poly-genotypes | 18/11/2016 | 37 | female | ERR12263835 |

### Culture adaptation, cloning, and high molecular weight DNA extraction

Since the isolates were derived from low-parasitaemia infections, parasites were culture-adapted to generate the high DNA yields necessary for long-read sequencing. Cloning by limiting dilution allowed the isolation of single-genotype populations (Supplementary Figure 1). Two extraction kits (Qiagen Genomic-tip 100/G and Monarch HMW DNA extraction kit) were tested for high molecular weight (HMW) DNA recovery, with the Monarch kit performing better (Supplementary Figure 2). Overall, a total set of 43 DNA extractions derived from 30 isolates were all successfully sequenced with PacBio HiFi (Table 1). Twenty three (23) genomes were derived from cloning dilutions and were strictly single-genotype parasite populations. The other 20 DNA samples were bulk populations (not cloned) with an unknown number of parasite genotypes. Based on their level of heterozygosity over a set of 72,020 informative loci, we inferred 8 bulk genomes to be single-genotype and 12 to consist of multiple parasite genotypes (Table 1). Across infection groups, we sequenced 17 genomes from asymptomatic infections in the dry season (DC group), 15 genomes from symptomatic infections in the wet season (WS group) and 11 genomes from asymptomatic infections in the wet season (WA group).

### Pacbio HiFi enables chromosome-scale genome assemblies

Raw assemblies from the 43 Pacbio HiFi reads datasets contained between 17 and 185 contigs (Table 2), with a median number of 54. The N50 statistics in all genomes (range 1,198,427 to 1,693,536 bp, median = 1,571,854 bp) were indicative of the presence of large chromosome-size contigs, which was further supported by syntenic analyse comparing newly assembled genomes to the 3D7 reference genome (Figure 2). The number of complete telomere-to-telomere contigs was however systematically lower than the 14 expected *P. falciparum* chromosomes (range 2 to 9, median = 4), demonstrating difficulties in assembling hypervariable sub-telomeric regions and complete telomeric contigs.

**Figure 1:**
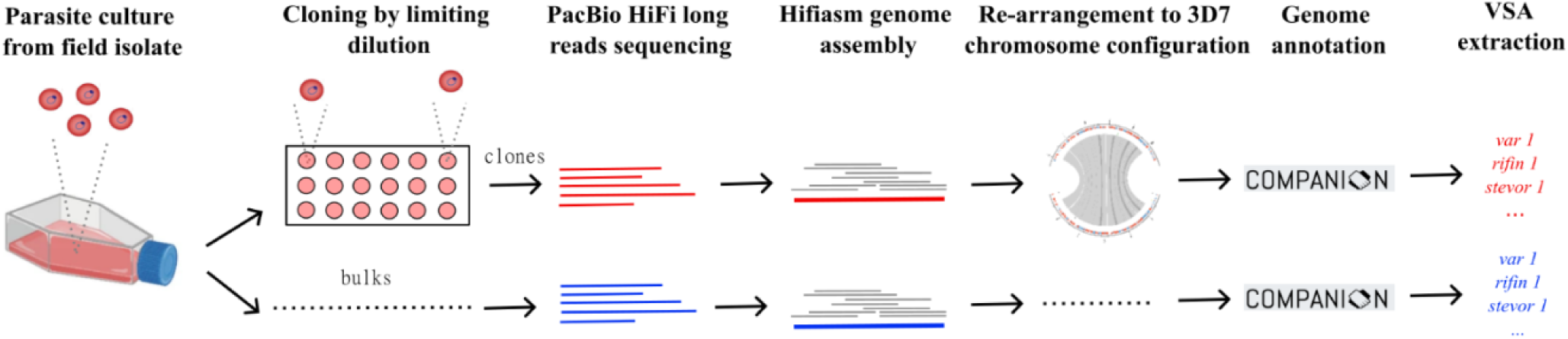
Schematic overview of the genome assembly method used to generate high quality genomes and extract Variant Surface Antigens (VSAs) from parasite cultures.

**Figure 2:**
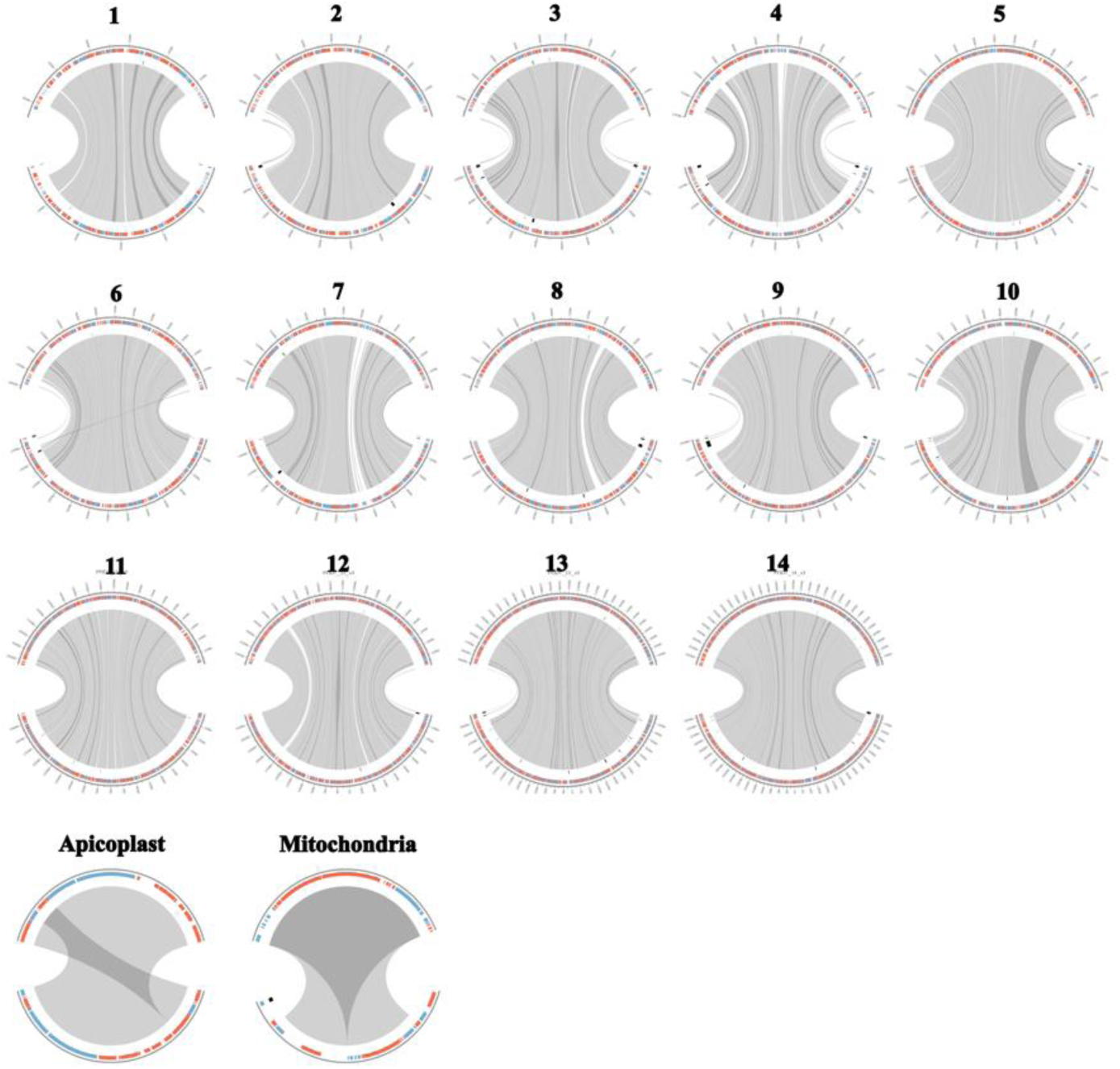
PacBio HiFi sequencing and hifiasm assembly allows chromosome-scale genome assemblies. Each circos plot represents chromosomes of the reference 3D7 genome (top) and the DC01m1 contigs (bottom) as a representative genome generated from single-genotype parasite lineage. Grey lines between top and bottom sequence assemblies indicate syntenic regions of high sequence similarity. The outer ring indicates position of genes annotated by the companion pipeline, genes in the forward strand are indicated in blue and genes in the reverse strand indicated in red. Gaps in synteny can be seen over internal *var* clusters on chromosomes 4, 7, 8 and 12.

**Table 2:** Summary statistics for 43 *P. falciparum* genome assemblies.

| ID | Genome length | Total contigs | Telomere to telomere contigs | N50 | L50 | Protein coding genes | var | LARSFADIG motifs | rif | stevor | var2csa |
| --- | --- | --- | --- | --- | --- | --- | --- | --- | --- | --- | --- |
| DC01m1c | 23.4 | 20 | 3 | 1622224 | 5 | 5718 | 70 | 61 | 107 | 24 | 2 |
| DC02m1c | 23.7 | 28 | 5 | 1693155 | 5 | 5751 | 70 | 64 | 178 | 34 | 1 |
| DC03m1c | 23.6 | 24 | 7 | 1627210 | 5 | 5760 | 68 | 64 | 156 | 34 | 1 |
| DC04m2c | 23.4 | 24 | 4 | 1636287 | 5 | 5721 | 66 | 62 | 118 | 25 | 2 |
| DC07m1b | 25.07 | 85 | 6 | 1433249 | 7 | 5473 | 73 | 70 | 172 | 35 | 4 |
| DC07m1c | 26.7 | 187 | 5 | 1507801 | 6 | 5532 | 60 | 58 | 154 | 34 | 2 |
| DC08m3b | 36.34 | 113 | 4 | 1198427 | 12 | 7970 | 116 | 115 | 230 | 43 | 4 |
| DC08m3c | 24.28 | 64 | 5 | 1631162 | 5 | 5388 | 55 | 55 | 134 | 32 | 2 |
| DC08m5b | 27.88 | 62 | 5 | 1421197 | 8 | 6192 | 105 | 103 | 234 | 47 | 4 |
| DC08m5c1 | 24.18 | 52 | 6 | 1606191 | 5 | 5428 | 73 | 71 | 133 | 22 | 2 |
| DC08m5c2 | 25 | 98 | 4 | 1630462 | 5 | 5478 | 62 | 62 | 150 | 32 | 2 |
| DC09m5b | 27.99 | 164 | 3 | 1405453 | 9 | 6003 | 116 | 114 | 236 | 46 | 2 |
| DC09m5c2 | 24.22 | 121 | 2 | 1661132 | 5 | 5489 | 84 | 79 | 130 | 28 | 1 |
| DC09m5c1 | 25.61 | 58 | 6 | 1543257 | 6 | 5482 | 67 | 66 | 161 | 33 | 1 |
| DC10m1c | 23.5 | 23 | 7 | 1556504 | 5 | 5378 | 70 | 68 | 134 | 26 | 1 |
| DC10m2c | 23.7 | 20 | 9 | 1571854 | 5 | 5415 | 71 | 69 | 154 | 29 | 2 |
| DC10m3c | 23.3 | 27 | 5 | 1672417 | 5 | 5384 | 52 | 63 | 137 | 23 | 2 |
| WA01b | 25.85 | 127 | 3 | 1423884 | 7 | 5411 | 65 | 64 | 116 | 25 | 2 |
| WA01c | 23.67 | 25 | 2 | 1637256 | 5 | 5383 | 69 | 67 | 120 | 25 | 2 |
| WA03b | 24.26 | 49 | 3 | 1519022 | 5 | 5513 | 72 | 69 | 164 | 37 | 3 |
| WA03c | 23.79 | 46 | 4 | 1693536 | 5 | 5465 | 70 | 69 | 154 | 36 | 6 |
| WA04b | 24.92 | 81 | 2 | 1446723 | 7 | 5456 | 75 | 73 | 165 | 35 | 4 |
| WA04c | 23.98 | 44 | 2 | 1613267 | 5 | 5519 | 83 | 76 | 170 | 37 | 4 |
| WA10b | 24.19 | 36 | 4 | 1487219 | 6 | 5416 | 82 | 80 | 162 | 35 | 5 |
| WA10c | 26.6 | 182 | 2 | 1468285 | 6 | 5539 | 79 | 77 | 161 | 33 | 5 |
| WA11c | 23.65 | 24 | 4 | 1621859 | 5 | 5399 | 73 | 70 | 105 | 25 | 2 |
| WA47b | 29.85 | 328 | 3 | 1408774 | 8 | 5568 | 91 | 86 | 177 | 39 | 9 |
| WA47c | 23.97 | 48 | 2 | 1693146 | 5 | 5481 | 71 | 66 | 160 | 36 | 4 |
| WS01b | 26.09 | 104 | 4 | 1517931 | 6 | 5759 | 124 | 119 | 193 | 39 | 1 |
| WS03b | 24.11 | 38 | 5 | 1658341 | 5 | 5487 | 79 | 76 | 132 | 33 | 4 |
| WS04b | 30.49 | 144 | 7 | 1440087 | 8 | 6682 | 131 | 130 | 293 | 54 | 5 |
| WS05b | 25.02 | 56 | 3 | 1435358 | 6 | 5615 | 72 | 70 | 166 | 45 | 1 |
| WS05c | 23.81 | 42 | 3 | 1641247 | 5 | 5506 | 68 | 66 | 153 | 42 | 1 |
| WS06c | 24.27 | 65 | 2 | 1560362 | 5 | 5466 | 64 | 62 | 149 | 34 | 4 |
| WS07c | 24.34 | 73 | 2 | 1612653 | 5 | 5492 | 69 | 66 | 158 | 37 | 2 |
| WS08b | 24.05 | 51 | 5 | 1638922 | 5 | 5428 | 67 | 66 | 116 | 25 | 2 |
| WS09b | 24.45 | 54 | 2 | 1653278 | 5 | 5518 | 65 | 64 | 163 | 35 | 1 |
| WS10b | 32.57 | 123 | 5 | 1386579 | 7 | 7432 | 87 | 85 | 165 | 39 | 1 |
| WS11b | 25.63 | 108 | 2 | 1479817 | 6 | 5887 | 69 | 66 | 152 | 33 | 1 |
| WS12b | 33.39 | 113 | 6 | 1356076 | 10 | 7222 | 130 | 130 | 274 | 51 | 2 |
| WS12c | 24.22 | 45 | 2 | 1649284 | 5 | 5498 | 75 | 75 | 152 | 27 | 1 |
| WS13b | 24.09 | 40 | 5 | 1564570 | 5 | 5490 | 75 | 75 | 174 | 23 | 4 |
| WS15b | 24.3 | 41 | 5 | 1592718 | 5 | 5506 | 74 | 74 | 173 | 23 | 4 |

We observed clear differences in assembly metrics between poly-genotype and single-genotype assemblies. Poly-genotype assemblies exhibited a larger genome size (median=24.9Mbp, range: 24.09-36.34Mbp), higher number of contigs (median = 118; range: 62–328) and protein-coding genes (median = 5881; range: 5487–7970). In contrast, single-genotype assemblies showed smaller genome size (median=24.19Mbp, range: 23.3-29.9Mbp), fewer contigs (median = 48; range: 20–187) and fewer protein-coding genes (median = 5481; range: 5388–5760). This difference was further confirmed in the number of VSAs, with poly-genotype assemblies harbouring more *var* genes (median = 118, range: 62-328) than their single-genotype counterpart (median = 70, range: 52-84).

Across the 31 single-genotype genomes, some of the Hifiasm assemblies contained contigs with regions syntenic to more than one chromosome. Mapping PacBio HiFi reads onto their assemblies revealed that the apparent chromosome breaks coincided with regions of low coverage or low sequence complexity, with no evidence of breaks supported by high read depth. Accordingly, raw contigs were split and reorganized into the 3D7 chromosome structure using ragtag (Alonge et al., 2022). After the re-contiguation step, all single-genotype assemblies had 14 large contigs corresponding to the 14 *P. falciparum* chromosomes, alongside 2 contigs for the mitochondrial and apicoplast genomes and a variable number of small extra contigs (median = 32, range 4 to 171; Figure 2, Table 2).

### Pacbio Hifi P. falciparum genome assemblies showed high base-calling accuracy and completeness

To assess base-calling accuracy of genome assemblies, we aligned Illumina reads from two ex vivo isolates, DC01m1 and DC10m2, (Guery et al., 2025) to their respective Pacbio HiFi assemblies from clonal genomes. We found only 55 SNPs over the entire DC01m1 genome (total length 23.6 Mbp) and 40 SNPs over the entire DC10m2 genome (total length 23.5 Mbp). We also computed k-mer-based quality scores using Merqury (Rhie et al., 2020), which yielded quality value (qv) scores of 64.5 for DC01m1 and 65.5 for DC10m2, indicating error rates ranging between 1 in 1,000,000 bp and 1 in 10,000,000 bp for both genomes. Completeness - the fraction of the expected genome sequenced found in an assembly-was equal to 94.6% in DC01m1 and 98.7% in DC10m2. Those results confirmed the high accuracy and quality of genome assemblies generated by hifiasm for Pacbio HiFi reads, and their use in generating reliable *P. falciparum* genome assemblies.

### IBD analysis reveals genotypic relatedness

To investigate genetic relatedness, we calculated identity-by-descent (IBD) fractions using core-genome SNPs mapped to the 3D7 reference genome (Supplementary Figure 3). In some cases, the same parasite genotypes were observed in different hosts (e.g. DC03m1c and WS11b), highlighting the presence and persistence of certain parasite genotypes across infection groups and time. In addition to shared parasite genotypes, we also observed groups of parasites with intermediate pairwise IBD values ranging from 0.1 to 0.9, highlighting groups of inter-related parasites that share a common recent ancestry (*e.g.* DC07m1c and DC10m3c).

### Full VSAs repertoires retrieved from PacBio HiFi assemblies

We extracted 3,288 *var* genes from all genomes (1,504 unique sequences) using Companion annotations. Clonal assemblies contained 56–84 *var* genes per genome (median = 69), consistent with expectations for single-genotype isolates. Bulk assemblies had 65–130 *var* genes, suggesting up to two co-infecting strains in some samples. The number of extracted *var* genes strongly correlated with the number of protein-coding DNA sequences (CDS) containing the LARSFADIG PfEMP1 motif (p < 2e-16), consistent with earlier findings (Otto et al., 2019).

Using the information on shared genotypes across samples deduced from pairwise IBD values (Supplementary Figure 3), we defined an unbiased dataset of unique single-genotype isolates (N = 17). Within this dataset, the numbers and proportions of each *var* upstream groups were relatively stable (Figure 3 and Table 2). Group A *var* genes ranged from 8 to 16 genes per genome with a median number of 11 per genome, group B *var* genes ranged from 29 to 46 per genome with a median number of 37 genes per genome and group C *var* genes ranged from 8 to 19 per genome with a median number of 12 per genome. These *var* group proportions are similar to those observed in 3D7 (A =10, B = 38, C = 13). In genomes sequenced from poly-genotype isolates the proportions of groups A, B and C were similar to the ones observed in clones, with a larger overall number of *var* genes (Table 2). Some *var* genes could not be reliably classified into any upstream group. This included genes with very divergent upstream sequences and *var* genes sequences located in small contigs where their 500bp upstream sequence could not be isolated.

**Figure 3:**
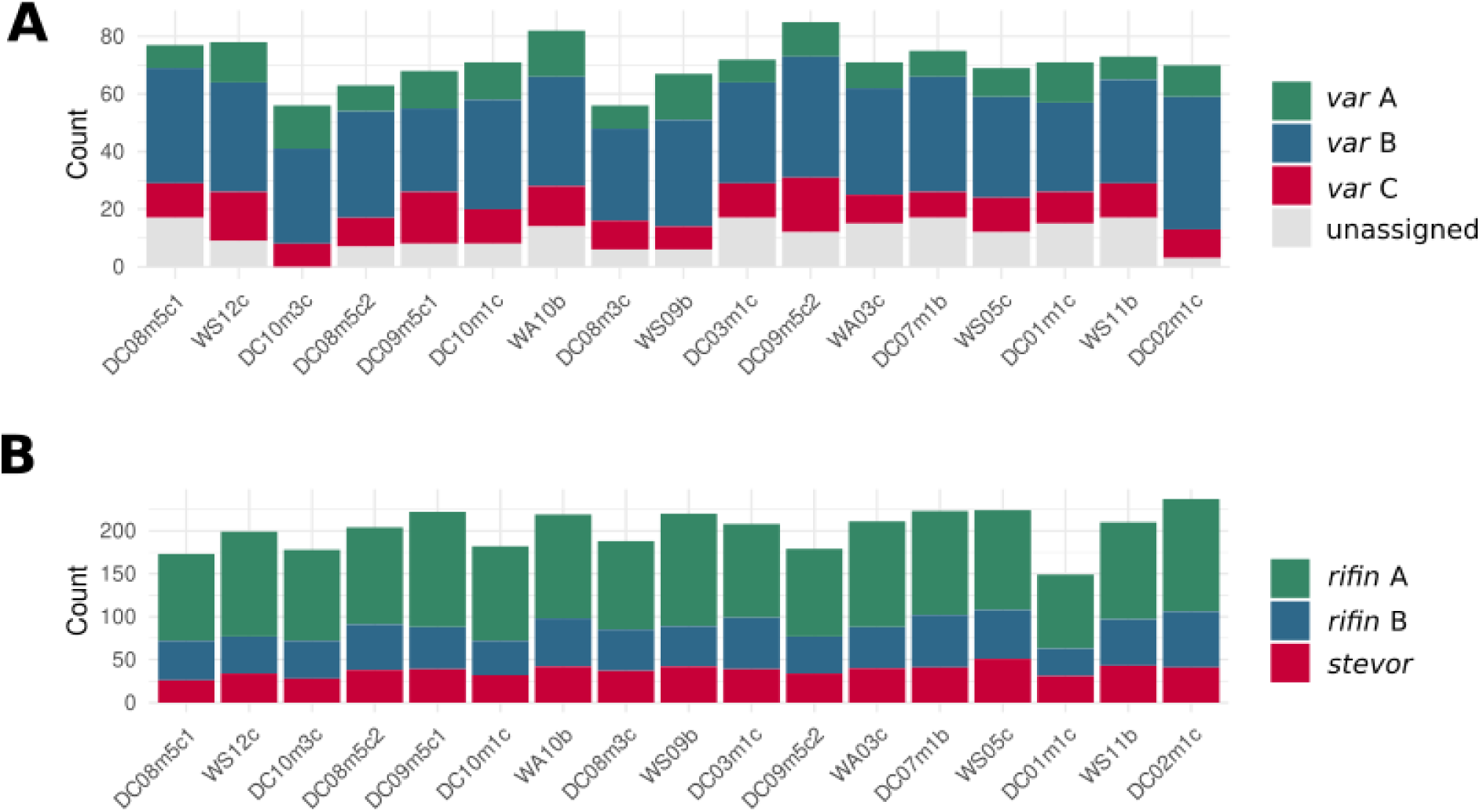
Numbers and proportions of VSA groups are stable across unique single-genotype assemblies (N = 17). The upper panel indicates the number of *var* gene per genome assembly, with colours indicating the different *var* groups. The lower panel indicates the number of *rif* and *stevor* per genome assembly, with colours indicating the different groups. Var genes for which the 500bp upstream was not available for classification were labelled ‘unassigned’.

As previously reported (Benavente et al., 2018; Sander et al., 2009), multiple copies of the conserved *var* gene *var2csa* were observed in multiple genomes (Table 2). Across the 17 unique single-genotype, 9 assemblies had a single *var2csa* copy and 8 assemblies had two or more *var2csa* copies (median = 2, range 1 to 3). In single-genotype genomes where copies of *var2csa* could be localised there was always a copy on chromosome 12, as in 3D7 and other reference genomes. Extra copies of the gene were found on sub-telomeric regions of chromosomes 1, 3, 8, 12, 13 and 14.

Similar to the pattern observed with *var* genes, the numbers and proportions of *rif* and *stevor* stayed relatively constant across different isolates (Figure 2). In the dataset of unique single-genotype genomes (N = 17) the total number of *rif* varied from 117 to 196 per genome with a median number of 168 genes. From those, *rif* group A ranged from 85 to 139 per genome with a median number of 116 genes and *rif* group B ranged from 32 to 65 with a median number of 48 genes per genome. These *rif* group proportions are similar to those observed in 3D7 (A = 111, B = 46). The *stevor* were less numerous, ranging from 26 to 51 with a median number of 39 genes per genome.

There was no correlation between the number of *var* genes and the number of *rif* (p = 0.61), or the number of *stevor* (p = 0.83), and the numbers of *rif* and *stevor* were mildly correlated (p = 0.03; see Supplementary Figure 4).

### Pairwise core genome IBD and VSAs sharing indexes are highly correlated

We quantified pairwise sharing indexes for *var*, *rif*, and *stevor* gene repertoires in the single-genotype dataset (N = 31, excluding technical replicates). The pairwise sharing indexes reflects a measure of genetic similarity for non-core *P. falciparum* genomes. Overall, sharing patterns closely paralleled core-genome IBD values—genome pairs with high IBD exhibited similar VSA repertoires (Figure 4A). Matrices permutation tests revealed significant correlations between core-genome IBD and VSA sharing for all three families (Figure 4B) (p < 0.01). Of note, we also observed most pairs of genomes seemed to share some *stevor* sequences, with an average *stevor* sharing value of 0.03 in pairs of samples with IBD < 0.25, against a lower average sharing value of 0.008 for *rif* and 0.003 for *var.* This is in line with previous observations that reported *stevor* as the least hypervariable of the three VSAs families (Q. Cheng et al., 1998; Lawton et al., 2025).

**Figure 4:**
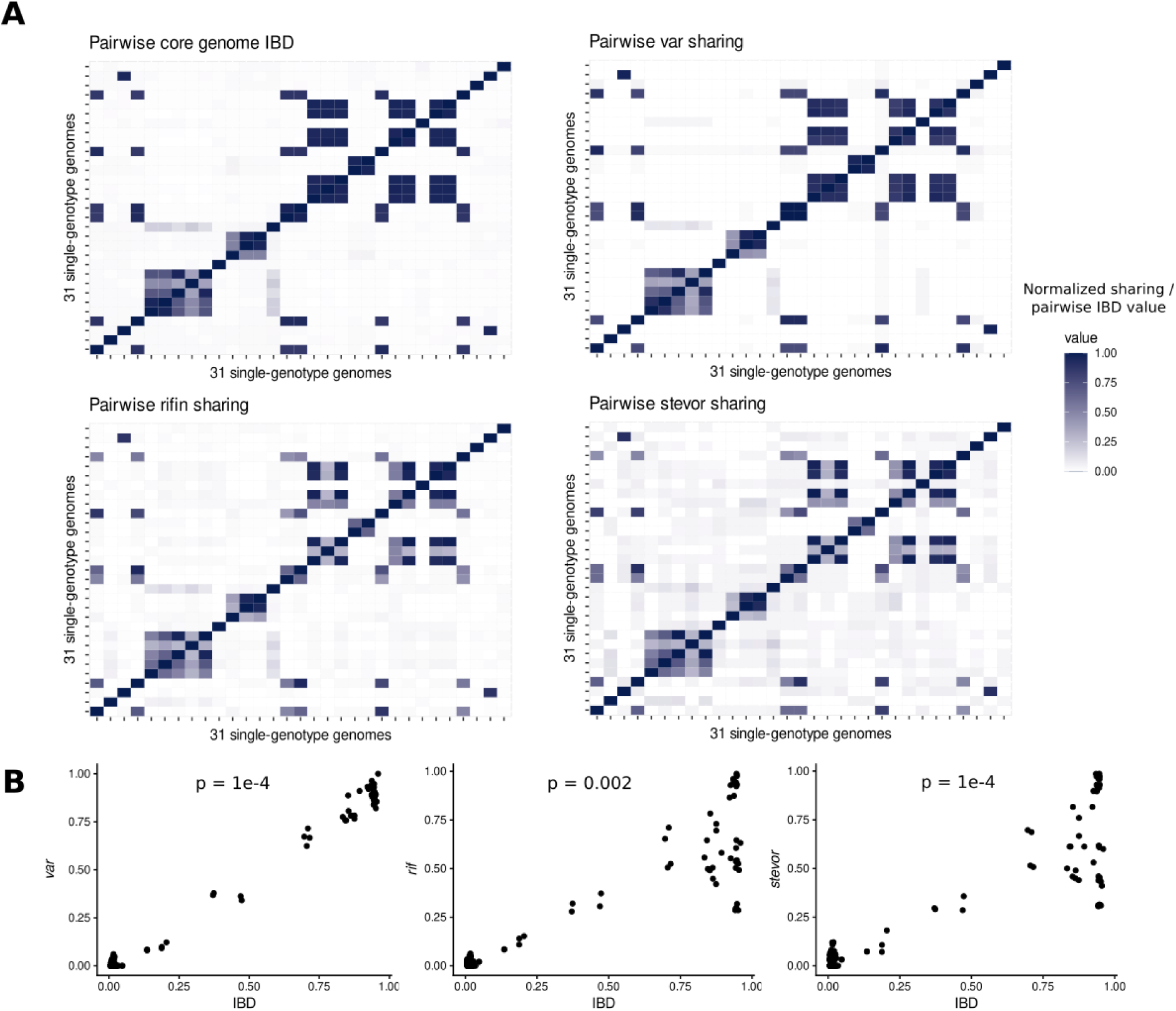
Groups of related parasites share common *var*, *rif* and *stevor* genes. (A) Pairwise matrices of IBD values and three normalized sharing indexes (*var*, *rif*, *stevor*) in a dataset of 31 single-genotype genomes (B) Correlations between pairwise IBD values and the three VSAs sharing indexes for the same dataset. P-values were calculated by a mantel test.

## Discussion

In total, we successfully culture-adapted and long-read sequenced parasite genomes from 43 parasite cultures, originating from 30 distinct field isolates. To the best of our knowledge this is the largest dataset of *P. falciparum* long read assemblies generated from natural infections to date. With PacBio HiFi reads and using the hifiasm assembler, we demonstrated long and accurate assembly contigs could be generated without the need for complementary illumina libraries, as is necessary with other long reads sequencing technologies (Delandre et al., 2024; Higgins et al., 2024). Particularly, our method proved useful in assembling *P. falciparum* sub-telomeric regions and complete sets of VSA families which are highly polymorphic and hard to assemble using short-read datasets (Andradi-Brown et al., 2024).

Culture adaptation was a necessary step to generate sufficient parasite DNA and to produce parasite lineages through cloning by limiting dilution. With this approach, parasite lineages could be delimited to ensure sequences were retrieved from single-genotype populations. Bulk populations (without the cloning step) also proved useful in retrieving complete VSA sequences, regardless of the number of parasite genotypes present.

The risk of culture-adaptation–induced mutations, although well established for parasites subjected to extended *in vitro* passages (Claessens et al., 2017, 2023; Kemp et al., 1992), is minimal here, as all genomes were derived from parasites maintained in culture only for the few weeks required for DNA extraction. This makes the generated assemblies a reliable genomic resource for analysis of circulating West African *P. falciparum*—across both symptomatic and asymptomatic infections. The alternative to culture adaptation is a Selective Whole Genome Amplification (SWGA) step using AT-rich primers and the phi29 polymerase. However, it remains to be tested whether this approach could replace the culture adaptation step without compromising on sequencing quality.

The analysis of VSA variation among genome assemblies confirmed patterns found in previous studies, with ratios of each *var* and *rif* groups stable across isolates. The presence of multiple copies of *var2csa* was observed in approximately half of the genomes generated here, which seems to be common in field isolates (Sander et al., 2009). There however seem to be an evolutionary constraint on maintaining at least one copy of the gene at the 5’ telomeric end of chromosome 12, as this copy was found in all genomes.

Contrary to a previous study which sampled *P. falciparum* reference genomes from widespread worldwide locations (Otto et al., 2018) we have generated a set of genomes originating from infections that were both close in collection dates and geographical locations. With this sampling design, we collected parasites with a wide range of genetic relatedness, from completely unrelated parasites to identical parasite genotypes sampled across different hosts or timepoints of a chronic infection. The relationship between core genome and non-core genome estimates of relatedness have seldom been studied (Neafsey et al., 2021), and the generation of genetic diversity in these two genomic regions stems from different biological processes. In the core genome, homologous recombination events occurring during meiosis lead to an uneven repartitioning of genetic variants and SNPs in offspring, with the proportion of the genome regions containing similar genetic variants tightly linked to the kinship value between parasites (Guo et al., 2025; Schaffner et al., 2018). In the non-core genome, variability is additionally generated through non-homologous recombination events, leading to sequence hypervariability and rapid generation of new VSAs (Claessens et al., 2014). Despite these differences, we observed a remarkably close relationship between core genome pairwise IBD values and the three sharing indexes for *var*, *rif*, *stevor*. These results indicate that in epidemiological settings similar to the one observed in The Gambia, measures of VSA sharing could effectively be used as a proxy for core genome relatedness, and vice-versa. These results support the validity of the growing number of studies relying on mass amplification of conserved *var* domains (e.g. DBLα) for population genetic analyses, and highlight VSA sharing as valid measure of genetic relatedness (Day et al., 2025).

The PacBio assemblies generated here additionally enable accurate detection of micro-indels, which are challenging to identify reliably using standard mapping of clinical isolates to the 3D7 reference genome. In a recent study, we used isolate-specific PacBio genomes as references for Illumina read mapping, allowing detection of micro-indels (Fogang et al., 2026). This approach allowed discrimination of parasite genomes from different patients that were otherwise indistinguishable using conventional Identity-By-Descent-based analyses. By combining PacBio assemblies with Illumina sequencing, we were able to track clonal parasite lineages in a setting of reduced malaria transmission. Such approaches are likely to become increasingly valuable in regions approaching malaria elimination, where ultra-fine-scale resolution of parasite transmission networks is required

Overall, the genome and VSA sequence datasets provided in this study presents a great resource for future research. Increasing the number of long read sequences generated from field isolated parasites will enable analyses of large structural variants occurring in *P. falciparum* genomes at a large scale, including regions with high repeats or high AT content. Furthermore, the complete and high-quality *var* gene sequences extracted here will prove a useful resource and training dataset for algorithm linking partial *var* sequences to full *var* domain composition (Mackenzie et al., 2022), or upstream group (Pangilinan et al., 2025). Finally, obtaining full-length VSA sequences will enable robust analyses of *var* gene expression in samples coming from The Gambia, and will enable differentiating *var* expression between asymptomatic and symptomatic infections.

## Data availability statement

All PacBio HiFi raw reads datasets are publicly available through the NCBI accession number indicated in Table 1. Assembled genomes are available on NCBI, BioProject PRJNA1368450. Annotation, VSA sequences and classification are available at the public repository https://zenodo.org/records/17551160. A series of codes and scripts used for this paper can be found on the github and zenodo repositories https://github.com/Mquenu/Falcigenomes-toolkit, https://zenodo.org/records/17724288.

## Acknowledgements

We thank Craig Corton for helpful advice on sample processing.

## References

Alonge, M., Lebeigle, L., Kirsche, M., Jenike, K., Ou, S., Aganezov, S., Wang, X., Lippman, Z. B., Schatz, M. C., & Soyk, S. (2022). Automated assembly scaffolding using RagTag elevates a new tomato system for high-throughput genome editing. Genome Biology, 23(1), 1–19. 10.1186/S13059-022-02823-7/FIGURES/2

Andradi-Brown, C., Wichers-Misterek, J. S., von Thien, H., Höppner, Y. D., Scholz, J. A. M., Hansson, H., Hocke, E. F., Gilberger, T. W., Duffy, M. F., Lavstsen, T., Baum, J., Otto, T. D., Cunnington, A. J., & Bachmann, A. (2024). A novel computational pipeline for var gene expression augments the discovery of changes in the Plasmodium falciparum transcriptome during transition from in vivo to short-term in vitro culture. ELife, 13. 10.7554/ELIFE.87726

Benavente, E. D., Oresegun, D. R., de Sessions, P. F., Walker, E. M., Roper, C., Dombrowski, J. G., de Souza, R. M., Marinho, C. R. F., Sutherland, C. J., Hibberd, M. L., Mohareb, F., Baker, D. A., Clark, T. G., & Campino, S. (2018). Global genetic diversity of var2csa in Plasmodium falciparum with implications for malaria in pregnancy and vaccine development. Scientific Reports 2018 8:1, 8(1), 1–8. 10.1038/s41598-018-33767-3

Cheng, H., Concepcion, G. T., Feng, X., Zhang, H., & Li, H. (2021). Haplotype-resolved de novo assembly using phased assembly graphs with hifiasm. Nature Methods 2021 18:2, 18(2), 170–175. 10.1038/s41592-020-01056-5

Cheng, Q., Cloonan, N., Fischer, K., Thompson, J., Waine, G., Lanzer, M., & Saul, A. (1998). stevor and rif are Plasmodium falciparum multicopy gene families which potentially encode variant antigens. Molecular and Biochemical Parasitology, 97(1–2), 161–176. 10.1016/S0166-6851(98)00144-3

Claessens, A., Adams, Y., Ghumra, A., Lindergard, G., Buchan, C. C., Andisi, C., Bull, P. C., Mok, S., Gupta, A. P., Wang, C. W., Turner, L., Arman, M., Raza, A., Bozdech, Z., & Rowe, J. A. (2012). A subset of group A-like var genes encodes the malaria parasite ligands for binding to human brain endothelial cells. Proceedings of the National Academy of Sciences of the United States of America, 109(26). 10.1073/PNAS.1120461109

Claessens, A., Affara, M., Assefa, S. A., Kwiatkowski, D. P., & Conway, D. J. (2017). Culture adaptation of malaria parasites selects for convergent loss-of-function mutants. Scientific Reports 2017 7:1, 7(1), 1–8. 10.1038/srep41303

Claessens, A., Hamilton, W. L., Kekre, M., Otto, T. D., Faizullabhoy, A., Rayner, J. C., & Kwiatkowski, D. (2014). Generation of Antigenic Diversity in Plasmodium falciparum by Structured Rearrangement of Var Genes During Mitosis. PLOS Genetics, 10(12), e1004812. 10.1371/JOURNAL.PGEN.1004812

Claessens, A., Stewart, L. B., Drury, E., Ahouidi, A. D., Amambua-Ngwa, A., Diakite, M., Kwiatkowski, D. P., Awandare, G. A., & Conway, D. J. (2023). Genomic variation during culture adaptation of genetically complex Plasmodium falciparum clinical isolates. Microbial Genomics, 9(5), 001009. 10.1099/MGEN.0.001009/CITE/REFWORKS

Collins, K. A., Ceesay, S., Drammeh, S., Jaiteh, F. K., Guery, M. A., Lanke, K., Grignard, L., Stone, W., Conway, D. J., D’Alessandro, U., Bousema, T., & Claessens, A. (2022). A Cohort Study on the Duration of Plasmodium falciparum Infections During the Dry Season in The Gambia. The Journal of Infectious Diseases, 226(1), 128–137. 10.1093/INFDIS/JIAC116

Danecek, P., Bonfield, J. K., Liddle, J., Marshall, J., Ohan, V., Pollard, M. O., Whitwham, A., Keane, T., McCarthy, S. A., & Davies, R. M. (2021). Twelve years of SAMtools and BCFtools. GigaScience, 10(2). 10.1093/GIGASCIENCE/GIAB008

Day, K. P., Tan, M. H., He, Q., Ruybal-Pesántez, S., Zhan, Q., Tiedje, K. E., & Pascual, M. (2025). Var genes, strain hyperdiversity, and malaria transmission dynamics. Trends in Parasitology, 41(6), 471–485. 10.1016/J.PT.2025.04.010

Delandre, O., Lamer, O., Loreau, J. M., Papa Mze, N., Fonta, I., Mosnier, J., Gomez, N., Javelle, E., & Pradines, B. (2024). Long-Read Sequencing and De Novo Genome Assembly Pipeline of Two Plasmodium falciparum Clones (Pf3D7, PfW2) Using Only the PromethION Sequencer from Oxford Nanopore Technologies without Whole-Genome Amplification. Biology, 13(2), 89. 10.3390/BIOLOGY13020089/S1

Fogang, B., Lellouche, L., Ceesay, S., Drammeh, S., Jaiteh, F. K., Guery, M. A., Landier, J., Haanappel, C. P., Froberg, J., Conway, D., D’Alessandro, U., Bousema, T., & Claessens, A. (2024). Asymptomatic Plasmodium falciparum carriage at the end of the dry season is associated with subsequent infection and clinical malaria in Eastern Gambia. Malaria Journal, 23(1). 10.1186/S12936-024-04836-Y

Fogang, B., Guery, M.-A., Cheeseman, I. H., Conway, D. J., & Claessens, A. (2026). Tracking malaria parasite lineages through de novo mutations in highly related Plasmodium falciparum genomes. 10.64898/2026.05.08.723589

Fu, L., Niu, B., Zhu, Z., Wu, S., & Li, W. (2012). CD-HIT: accelerated for clustering the next-generation sequencing data. Bioinformatics, 28(23), 3150. 10.1093/BIOINFORMATICS/BTS565

Gardner, M. J., Hall, N., Fung, E., White, O., Berriman, M., Hyman, R. W., Carlton, J. M., Pain, A., Nelson, K. E., Bowman, S., Paulsen, I. T., James, K., Eisen, J. A., Rutherford, K., Salzberg, S. L., Craig, A., Kyes, S., Chan, M. S., Nene, V., … Barrell, B. (2002). Genome sequence of the human malaria parasite Plasmodium falciparum. Nature, 419(6906), 498. 10.1038/nature01097

Gen, M., Ahouidi, A., Ali, M., Almagro-Garcia, J., Amambua-Ngwa, A., Amaratunga, C., Amato, R., Amenga-Etego, L., Andagalu, B., Anderson, T. J. C., Andrianaranjaka, V., Apinjoh, T., Ariani, C., Ashley, E. A., Auburn, S., Awandare, G. A., Ba, H., Baraka, V., Barry, A. E., … Ye, H. (2021). An open dataset of Plasmodium falciparum genome variation in 7,000 worldwide samples. Wellcome Open Research, 6, 42. 10.12688/WELLCOMEOPENRES.16168.2

Godin, M. J., Sebastian, A., Albert, I., & Lindner, S. E. (2023). Long-read genome assembly and gene model annotations for the rodent malaria parasite Plasmodium yoelii 17XNL. Journal of Biological Chemistry, 299(7). 10.1016/J.JBC.2023.104871/ATTACHMENT/BE8F4C96-D7A6-465F-A437-A435642F188E/MMC9.PDF

Guery, M.-A., Ceesay, S., Drammeh, S., Jaiteh, F. K., d’Alessandro, U., Bousema, T., Conway, D. J., & Claessens, A. (2025). Household clustering and seasonal genetic variation of Plasmodium falciparum at the community-level in The Gambia. ELife, 13. 10.7554/ELIFE.103047.2

Guo, B., Rowley, E., O’Connor, T. D., & Takala-Harrison, S. (2025). Potential and pitfalls of using identity-by-descent for malaria genomic surveillance. Trends in Parasitology, 41(5), 387–400. 10.1016/J.PT.2025.03.012/ATTACHMENT/04E8641E-AA9E-41CF-959F-32D78B225F93/MMC1.DOCX

Harrison, T. E., Mørch, A. M., Felce, J. H., Sakoguchi, A., Reid, A. J., Arase, H., Dustin, M. L., & Higgins, M. K. (2020). Structural basis for RIFIN-mediated activation of LILRB1 in malaria. Nature, 587(7833), 309–312. 10.1038/S41586-020-2530-3

Higgins, M., Manko, E., Ward, D., Phelan, J. E., Nolder, D., Sutherland, C. J., Clark, T. G., & Campino, S. (2024). New reference genomes to distinguish the sympatric malaria parasites, Plasmodium ovale curtisi and Plasmodium ovale wallikeri. Scientific Reports 2024 14:1, 14(1), 1–10. 10.1038/s41598-024-54382-5

Holzschuh, A., Lerch, A., Fakih, B. S., Aliy, S. M., Ali, M. H., Ali, M. A., Bruzzese, D. J., Yukich, J., Hetzel, M. W., & Koepfli, C. (2024). Using a mobile nanopore sequencing lab for end-to-end genomic surveillance of Plasmodium falciparum: A feasibility study. PLOS Global Public Health, *4*(2 February), e0002743. 10.1371/JOURNAL.PGPH.0002743/OG_IMAGE.JPG

Hon, T., Mars, K., Young, G., Tsai, Y. C., Karalius, J. W., Landolin, J. M., Maurer, N., Kudrna, D., Hardigan, M. A., Steiner, C. C., Knapp, S. J., Ware, D., Shapiro, B., Peluso, P., & Rank, D. R. (2020). Highly accurate long-read HiFi sequencing data for five complex genomes. Scientific Data 2020 7:1, 7(1), 1–11. 10.1038/s41597-020-00743-4

Jespersen, J. S., Wang, C. W., Mkumbaye, S. I., Minja, D. T., Petersen, B., Turner, L., Petersen, J. E., Lusingu, J. P., Theander, T. G., & Lavstsen, T. (2016). Plasmodium falciparum var genes expressed in children with severe malaria encode CIDRα1 domains. EMBO Molecular Medicine, 8(8), 839–850. 10.15252/EMMM.201606188

Kemp, D. J., Thompson, J., Barnes, D. A., Triglia, T., Karamalis, F., Petersen, C., Brown, G. V., & Day, K. P. (1992). A chromosome 9 deletion in Plasmodium falciparum results in loss of cytoadherence. Memorias Do Instituto Oswaldo Cruz, 87 *Suppl 3*, 85–89. 10.1590/S0074-02761992000700011

Knaus, B. J., & Grünwald, N. J. (2017). vcfr: a package to manipulate and visualize variant call format data in R. Molecular Ecology Resources, 17(1), 44–53. 10.1111/1755-0998.12549

Lavstsen, T., Salanti, A., Jensen, A. T., Arnot, D. E., & Theander, T. G. (2003). Sub-grouping of Plasmodium falciparum 3D7 var genes based on sequence analysis of coding and non-coding regions. Malaria Journal, 2(1), 27. 10.1186/1475-2875-2-27

Lawton, J. G., Zhou, A. E., Stucke, E. M., Takala-Harrison, S., Silva, J. C., & Travassos, M. A. (2025). Diamonds in the rif: Alignment-free comparative genomics analysis reveals strain-transcendent Plasmodium falciparum antigens amidst extensive genetic diversity. Infection, Genetics and Evolution, 129, 105725. 10.1016/J.MEEGID.2025.105725

Li, H. (2018). Minimap2: pairwise alignment for nucleotide sequences. Bioinformatics, 34(18), 3094–3100. 10.1093/BIOINFORMATICS/BTY191

Li, H., & Durbin, R. (2009). Fast and accurate short read alignment with Burrows–Wheeler transform. Bioinformatics, 25(14), 1754. 10.1093/BIOINFORMATICS/BTP324

Li, H., Handsaker, B., Wysoker, A., Fennell, T., Ruan, J., Homer, N., Marth, G., Abecasis, G., & Durbin, R. (2009). The Sequence Alignment/Map format and SAMtools. Bioinformatics, 25(16), 2078–2079. 10.1093/BIOINFORMATICS/BTP352

Mackenzie, G., Jensen, R. W., Lavstsen, T., & Otto, T. D. (2022). Varia: a tool for prediction, analysis and visualisation of variable genes. BMC Bioinformatics, 23(1). 10.1186/S12859-022-04573-6

Neafsey, D. E., Taylor, A. R., & MacInnis, B. L. (2021). Advances and opportunities in malaria population genomics. Nature Reviews. Genetics, 22(8), 502. 10.1038/S41576-021-00349-5

Otto, T. D., Assefa, S. A., Böhme, U., Sanders, M. J., Kwiatkowski, D. P., Berriman, M., & Newbold, C. (2019). Evolutionary analysis of the most polymorphic gene family in Falciparum malaria. Wellcome Open Research, 4. 10.12688/WELLCOMEOPENRES.15590.1/DOI

Otto, T. D., Böhme, U., Sanders, M., Reid, A., Bruske, E. I., Duffy, C. W., Bull, P. C., Pearson, R. D., Abdi, A., Dimonte, S., Stewart, L. B., Campino, S., Kekre, M., Hamilton, W. L., Claessens, A., Volkman, S. K., Ndiaye, D., Amambua-Ngwa, A., Diakite, M., … Berriman, M. (2018). Long read assemblies of geographically dispersed Plasmodium falciparum isolates reveal highly structured subtelomeres. Wellcome Open Research, 3, 52. 10.12688/WELLCOMEOPENRES.14571.1

Otto, T. D., Dillon, G. P., Degrave, W. S., & Berriman, M. (2011). RATT: Rapid Annotation Transfer Tool. Nucleic Acids Research, 39(9), e57–e57. 10.1093/NAR/GKQ1268

Pangilinan, E. A., Quenu, M., Claessens, A., & Otto, T. D. (2025). upsAI: A high-accuracy machine learning classifier for predicting Plasmodium falciparum var gene upstream groups. BioRxiv, 2025.05.19.654848. 10.1101/2025.05.19.654848

Rask, T. S., Hansen, D. A., Theander, T. G., Pedersen, A. G., & Lavstsen, T. (2010). Plasmodium falciparum erythrocyte membrane protein 1 diversity in seven genomes--divide and conquer. PLoS Computational Biology, 6(9). 10.1371/JOURNAL.PCBI.1000933

Rhie, A., Walenz, B. P., Koren, S., & Phillippy, A. M. (2020). Merqury: Reference-free quality, completeness, and phasing assessment for genome assemblies. Genome Biology, 21(1), 1–27. 10.1186/S13059-020-02134-9/FIGURES/6

Rowe, J. A., Claessens, A., Corrigan, R. A., & Arman, M. (2009). Adhesion of Plasmodium falciparum-infected erythrocytes to human cells: Molecular mechanisms and therapeutic implications. Expert Reviews in Molecular Medicine, 11. 10.1017/S1462399409001082

Sakoguchi, A., Chamberlain, S. G., Mørch, A. M., Widdess, M., Harrison, T. E., Dustin, M. L., Arase, H., Higgins, M. K., & Iwanaga, S. (2025). RIFINs displayed on malaria-infected erythrocytes bind KIR2DL1 and KIR2DS1. Nature 2025, 1–9. 10.1038/s41586-025-09091-y

Sander, A. F., Salanti, A., Lavstsen, T., Nielsen, M. A., Magistrado, P., Lusingu, J., Ndam, N. T., & Arnot, D. E. (2009). Multiple var2csa-Type PfEMP1 Genes Located at Different Chromosomal Loci Occur in Many Plasmodium falciparum Isolates. PLOS ONE, 4(8), e6667. 10.1371/JOURNAL.PONE.0006667

Schaffner, S. F., Taylor, A. R., Wong, W., Wirth, D. F., & Neafsey, D. E. (2018). HmmIBD: Software to infer pairwise identity by descent between haploid genotypes. Malaria Journal, 17(1), 1–4. 10.1186/S12936-018-2349-7/FIGURES/1

Stanke, M., Steinkamp, R., Waack, S., & Morgenstern, B. (2004). AUGUSTUS: a web server for gene finding in eukaryotes. Nucleic Acids Research, 32(Web Server issue), W309. 10.1093/NAR/GKH379

Tonkin-Hill, G., Ruybal-Pesántez, S., Tiedje, K. E., Rougeron, V., Duffy, M. F., Zakeri, S., Pumpaibool, T., Harnyuttanakorn, P., Branch, O. L. H., Ruiz-Mesía, L., Rask, T. S., Prugnolle, F., Papenfuss, A. T., Chan, Y. B., & Day, K. P. (2021). Evolutionary analyses of the major variant surface antigen-encoding genes reveal population structure of Plasmodium falciparum within and between continents. PLOS Genetics, 17(2), e1009269. 10.1371/JOURNAL.PGEN.1009269

Van der Auwera, G., O’Connor, B., & Safari. (2020). Genomics in the Cloud: Using Docker, GATK, and WDL in Terra. O’Reilly Media, 300. https://www.oreilly.com/library/view/genomics-in-the/9781491975183/

Vembar, S. S., Seetin, M., Lambert, C., Nattestad, M., Schatz, M. C., Baybayan, P., Scherf, A., & Smith, M. L. (2016). Complete telomere-to-telomere de novo assembly of the Plasmodium falciparum genome through long-read (>11 kb), single molecule, real-time sequencing. DNA Research, 23(4), 339–351. 10.1093/DNARES/DSW022

Wenger, A. M., Peluso, P., Rowell, W. J., Chang, P. C., Hall, R. J., Concepcion, G. T., Ebler, J., Fungtammasan, A., Kolesnikov, A., Olson, N. D., Töpfer, A., Alonge, M., Mahmoud, M., Qian, Y., Chin, C. S., Phillippy, A. M., Schatz, M. C., Myers, G., DePristo, M. A., … Hunkapiller, M. W. (2019). Accurate circular consensus long-read sequencing improves variant detection and assembly of a human genome. Nature Biotechnology 2019 37:10, 37(10), 1155–1162. 10.1038/s41587-019-0217-9

World malaria report 2024. (n.d.). Retrieved July 25, 2025, from https://www.who.int/teams/global-malaria-programme/reports/world-malaria-report-2024

Zhou, A. E., Shah, Z. V., Bradwell, K. R., Munro, J. B., Berry, A. A., Serre, D., Takala-Harrison, S., O’Connor, T. D., Silva, J. C., & Travassos, M. A. (2022). STRIDE: a command-line HMM-based identifier and sub-classifier of Plasmodium falciparum RIFIN and STEVOR variant surface antigen families. BMC Bioinformatics, 23(1), 1–9. 10.1186/S12859-021-04515-8/TABLES/2

