## Supplementary files for "De novo assembly of complete *Plasmodium falciparum* isolate genomes using PacBio HiFi sequencing technology"

### Supplementary methods

*DNA extraction and parasite genotyping with MSP 1 and 2*

Fifty microlitre (50µL) aliquots of thawed or culture adapted sample were taken for DNA extracted. Extraction was done with the NucleoSpin Blood QuickPure kit (Macherey-Nagel), following manufacturer's protocol. DNA was eluted with 20µL of nuclease-free water. Clonality of each sample was assessed by genotyping the highly polymorphic merozoite surface proteins 1 and 2 (MSP1 and MSP2), using previous described primers (Somé *et al.*, 2018). The repetitive block 2 of MSP1 and block 3 of MSP2 (Smythe *et al.*, 1990) were amplified with conserved sequence primers, followed by a nested step where the K1, MAD20 and R033 alleles of MSP1 and the IC/3D7 and FC27 alleles of MSP2 were amplified with allele-specific primers. First Polymerase Chain Reaction (PCR) was performed using 5µL genomic DNA and the GoTaq Green Master Mix (Promega). Cycling conditions were as follows: initial denaturation at 95°C for 5 minutes, followed by 30 cycles of denaturation at 94°C for 1 minute, annealing at 50°C for 45 seconds and extension at 72°C for 1.5 minutes, with final extension of 72°C for 10 minutes. PCR products were diluted ten-fold and 1µL used as template for the nested PCR with the following conditions: initial denaturation at 95°C for 5 minutes, followed by 25 cycles of denaturation at 94°C for 45 seconds, annealing at 61°C for 45 seconds and extension at 72°C for 45 seconds, with final extension of 72°C for 5 minutes. Primers were used at a concentration of 200ng for both assays. PCR products were resolved on 2% ethidium bromide-stained agarose gels.

### Supplementary results

*Comparison of the two extraction kits Qiagen and Monarch*

DNA extraction trials on a subset of cultures were performed with the Qiagen genomic-tip 100/G kit and Monarch HMW DNA Extraction Kit for Tissue. Both extraction methods were capable of recovering high molecular weight genomic DNA larger than 10 kilobase pairs (kb) as determined by 0.5% agarose gel electrophoresis (Supplementary Figure 2). Size analysis with the Femto Pulse System (Agilent) showed larger DNA size fragments ($P$ = 0.002; t-test) in the extracts from the Monarch kit (ranging from 72kb to 183kb – 229kb on average), compared to the Qiagen kit (43kbp to 96kb – 62kb on average, Supplementary Figure 2). The Monarch extraction kit was therefore used for all DNA extractions sent for sequencing.

### Supplementary Figures

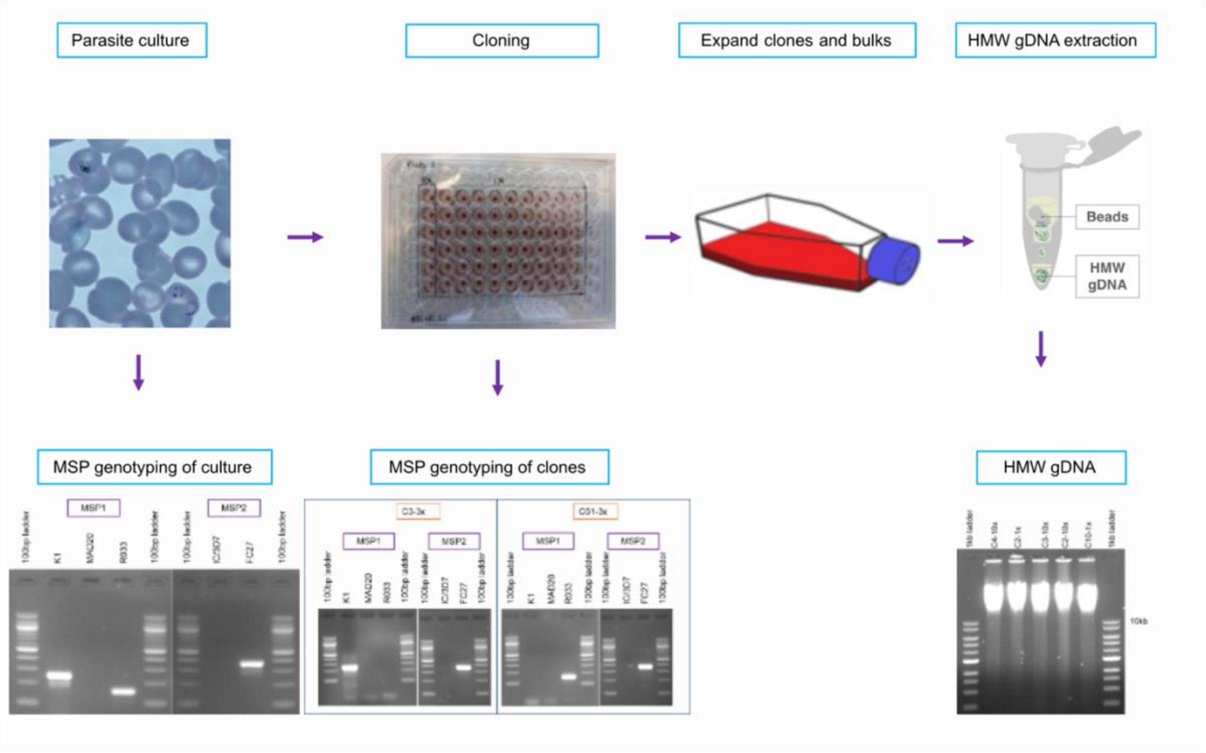


**Supplementary Figure 1:** Overall design of the lab procedure to generate High Molecular Weight DNA extracts from *Plasmodium falciparum* field isolates.

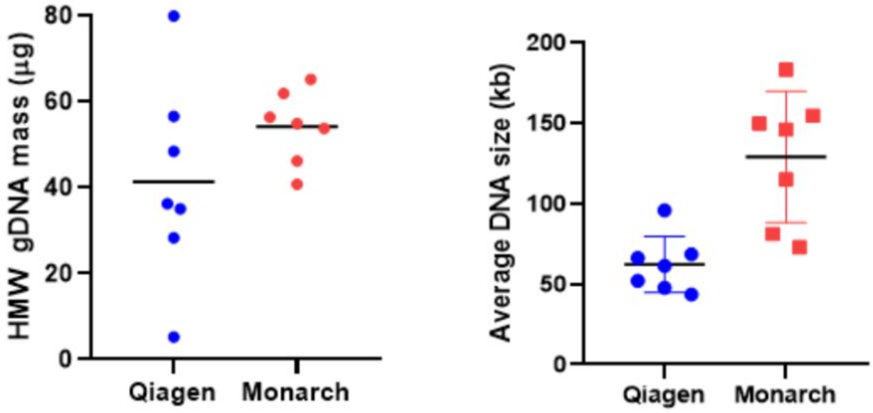


**Supplementary Figure 2:** Comparison of the two extraction kits Qiagen and Monarch for high molecular weight DNA extraction.

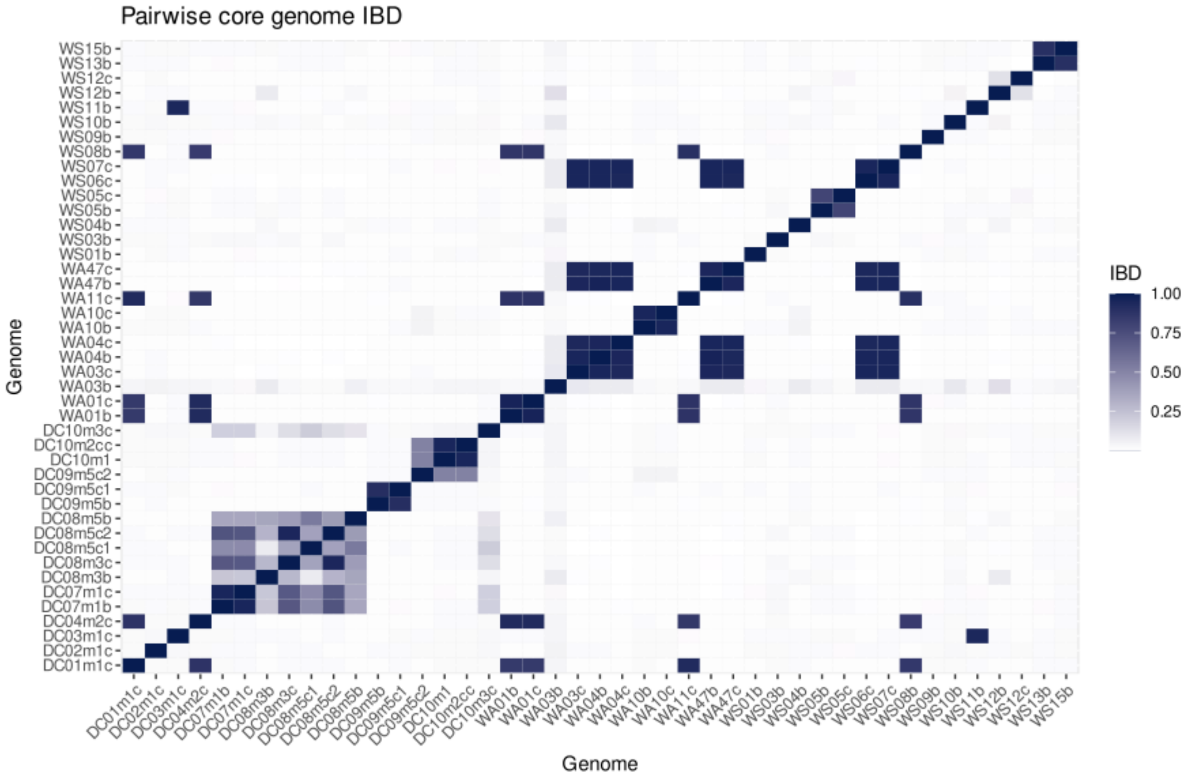


**Supplementary Figure 3: Pairwise IBD values (proportion of chromosome segment in IBD state) between all of the 43 *P. falciparum* genome assemblies.**

As expected, genomes from both clone and single-genotype bulk populations from the same field isolate had pairwise IBD values > 0.9, confirming those samples were effectively technical replicates of the same field isolate parasite genome (*e.g.* WA01b and WA01c). Similarly, we sometimes sampled the same parasite genotypes at different timepoints within a chronic infection (*e.g.* DC08m3c and DC08m5c2).

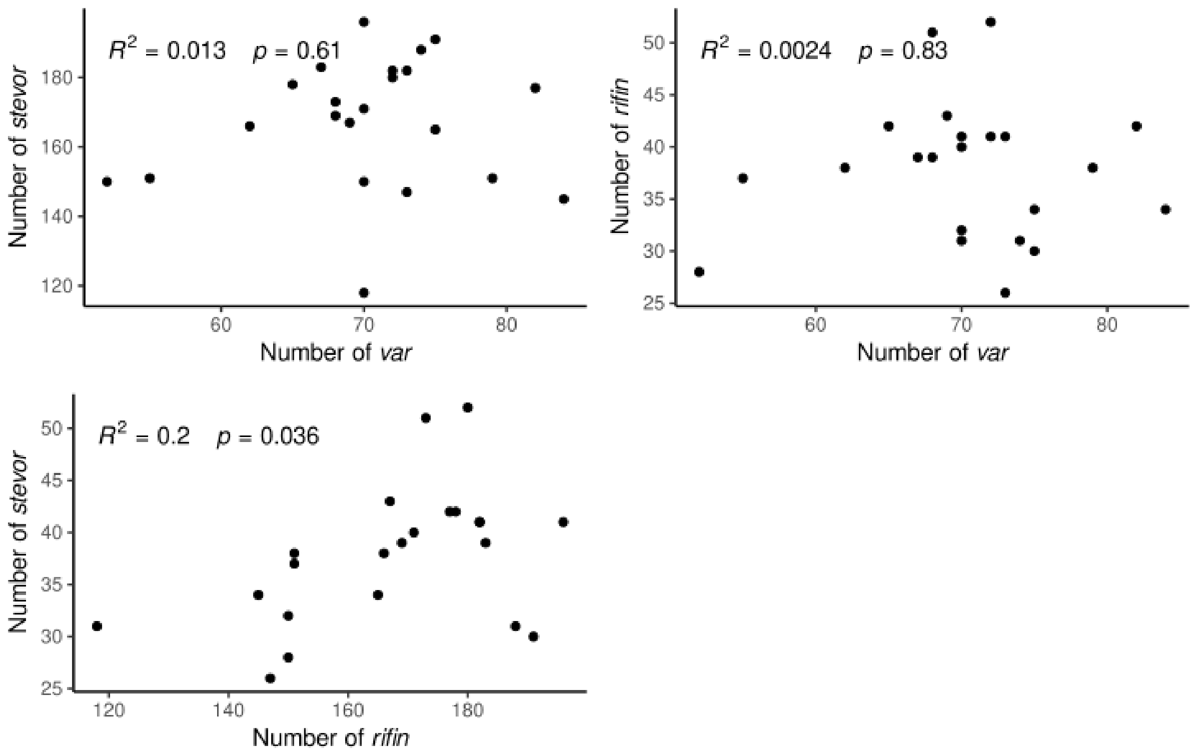


**Supplementary Figure 4:** Correlations between the number of VSAs across unique single-genotype genomes (N = 22).
